# Test-Retest Reliability Analysis of Resting-state EEG Measures and Their Association with Long-Term Memory in Children and Adults

**DOI:** 10.1101/2025.08.04.668389

**Authors:** Anastasios Ziogas, Simon Ruch, Nicole H. Skieresz, Sandy C. Marca, Nicolas Rothen, Thomas P. Reber

## Abstract

EEG resting-state measures, such as spectral power and microstates, have been associated with human long-term memory (LTM) performance. However, findings across studies are inconsistent and sometimes contradictory, likely due to a low reliability of the measures employed. These inconsistencies limit the interpretability and generalizability of results, emphasizing the need for a systematic evaluation of measure reliability. In this study, we addressed this gap by identifying the most reliable EEG resting-state measures and evaluating their predictive value for LTM performance in a second-language (L2) vocabulary learning paradigm. A group of children (N = 36) and adults (N = 90) participated in 2 studies on app-assisted learning of second language vocabulary. Participants completed a test on L2 vocabulary and a resting-state EEG recording (180 s eyes open) before and after learning a second language using a smartphone app. We used Intraclass Correlation Coefficients (ICC) to identify resting state EEG measures with satisfying test-retest reliability (ICC >= 0.75) and then assessed how these reliable measures are associated with L2 vocabulary learning representing LTM performance. Highest ICC values were found for oscillatory power in the alpha range and in the frequency of occurrences, duration and coverages of microstates. Calculations yielded ICC values of 0.84/0.86 (children/adults) for alpha power and 0.88/0.80 for microstate measures. Of these measures, only alpha power showed a positive correlation with LTM performance, but only in the adult population (*r* = 0.38, *p* < .01). No other measures were associated with LTM (all *p* > 0.05). Alpha power could thus serve as a stable and reliable marker of the neural mechanisms accounting for high LTM performance in the fully developed adult brain.

## 1 Introduction

Our understanding of the neurobiology of human long-term-memory (LTM) is consequential for our conceptualization of the uniqueness of human cognition. Prominent classical cognitive models have often implemented some form of language memory in their framework (Baddeley, 2003) as memory is essential for the development of language (Corballis, 2019). This even led to assumptions on an independently evolved link between verbal working memory and verbal LTM (Hartshorne & Makovski, 2019) further promoted by evidence that LTM is supported by the same neural systems involved in language comprehension and production (Schwering & MacDonald, 2020). Language stimuli or language skills over time represent a useful tool to study LTM with its neurobiological components and EEG is a useful method to capture the temporal dynamics of language processes and their association to LTM. In EEG resting-state recordings, the participants are not required to perform a task or perceive any stimulation. The minimal demands for participants make it easier to study e.g., children. Still, this line of research has provided heterogeneous findings on LTM or language within different developmental stages (Amin & Malik, 2013; Gaudet et al., 2020; Khader & Rösler, 2011; Lui et al., 2021). This could be partly due to the multitude of methodological approaches available to handle resting state EEG data e.g., calculations of EEG power, microstates.

The most frequently used measures and parameters in the field of resting-state EEG are the absolute or relative oscillatory power for the prominent frequency ranges (e.g., delta: 0.5-4 Hz). Oscillatory power has often been linked directly to LTM (Brokaw et al., 2016; Murphy et al., 2018), long-term potentiation (Savarimuthu & Ponniah, 2024) and memory consolidation, also when recorded intracranially (Axmacher et al., 2008) or when different stimulus modalities were used (e.g., auditory LTM, Lenz et al., 2007). EEG power measures have been shown to be sensitive to both LTM encoding and retrieval (Babiloni et al., 2004). In a recent study EEG power was used to successfully predict semantic LTM formation after two, four and six months, outperforming other behavioral based memory assessments (e.g., Raven’s Advanced Progressive Matrices; Amin et al., 2025). Especially alpha power has been found to correlate with LTM performance (Khader & Rösler, 2011) and activation of semantic concepts (Klimesch, et al., 1997a; Klimesch et al., 1997b). It is assumed that brain activity in the alpha band represents the inhibition of irrelevant mental processes, facilitating memory formation and regulating the flow of information through the brain (Jensen & Mazaheri, 2010). A decrease in alpha power was also demonstrated through sleep deprivation (24 and 36 hours) (Lian et al., 2023) and sleep has been suggested as an essential process for LTM formation (Klinzing et al., 2019).

Early works on EEG power and language have shown lower theta (Colon et al., 1979; Sklar et al., 1972) and higher alpha power (Duffy et al., 1980) in small samples of healthy children compared to dyslexic children. More significant changes in these frequency bands were observed over a period of 2.5–3 years in poor readers compared to typical readers (Harmony et al., 1995). This was interpreted as evidence of maturational delay in readers with impairments. The findings in the theta range were replicated in later studies on children (Arns et al., 2007) and also in adult dyslexics (Rumsey et al., 1989). Low theta power localized in the left hemisphere was also related to higher sentence comprehension (Beese et al., 2017). The pattern of diminished low frequency (delta, theta) power and pronounced higher frequency (alpha, beta, gamma) power was often related to better language function throughout the literature (Barry et al., 2009; Benasich et al., 2008; Brito et al., 2016; Clarke et al., 2002; Gou et al., 2011; Lum et al., 2022; Papagiannopoulou & Lagopoulos, 2016; Tierney et al., 2014) while some studies did not replicate this pattern (Babiloni et al., 2012; Lui et al., 2021; Nazari et al., 2012; Pinkerton et al., 1989; Schiavone et al., 2014; Shiota et al., 2000). When EEG-neurofeedback was used to train a group of children with reading disabilities, improvement in reading was not related to any changes in resting-state-EEG power (Nazari et al., 2012). While it remains difficult to integrate all these findings, this simple calculation of power values on the scalp provides by far the largest amount of empirical data on resting state EEG measures associated with LTM or language functions over time in the human brain.

Less common is the calculation of so-called microstates or estimations about the cortical sources of EEG signals generated on the scalp. Microstate analyses have the advantage that they rely on multivariate data-driven methods (Michel & Koenig, 2018) that do not require predefining specific frequencies, time windows, or scalp regions of interest. Microstates are unique electric fields or voltage topographies in the scalp EEG that repeatedly occur for brief, quasi-stable periods (Lehmann et al., 1987; Strik & Lehmann, 1993). Different states are thought to represent distinct neuronal generators and mechanisms in the brain (Lehmann et al., 1987; Strik & Lehmann, 1993). Microstates also rely on relatively simple computation methods (Khanna et al., 2015) producing reliable results (Kleinert et al., 2023) and have been linked directly to memory consolidation of word-pairs (Poskanzer et al., 2021). Usually, a set of prototypical (e.g., four) microstates are identified in the recorded segment and used to label the entire resting state recording into sequences of the prototypical microstates. Several statistics (e.g., Global Field Power, frequency of occurrence) can then be estimated for every microstate and used for further analysis. When looking at microstates at the scalp, one study reported a negative correlation between the microstate class E and a language test (Mini Mental State Examination subtest) (Grieder et al., 2016). Right-lateralized sources of microstate class B were related to verbal processing in another study (Milz et al., 2016).

Some studies have analyzed the reliability of EEG power and microstates (e.g., Lopez et al., 2023; Popov et al., 2023); however, these measures have not yet been directly linked to LTM performance. The aim of this study is to explore the reliability of EEG power and microstate measures at the scalp level and their relation to LTM performance while accounting for developmental effects by including different age groups. High split-half and test-retest reliability are essential prerequisites for using a measure as a potential marker of the neural architecture and processes underlying LTM performance, as they ensure the consistency and stability necessary for meaningful interpretation. Within a longitudinal study on second language acquisition using an educational app for Swiss schoolchildren and adults (Reber & Rothen, 2018), the experimental design already provides an optimal opportunity to quantify split-half and test-retest reliability for the candidate resting state measures and directly relate them to language improvement. This study involves participants of both age groups completing a resting state EEG session and a language test at two timepoints, in between which they used an app to learn vocabulary of a second language. Using this setting, the collected data at both timepoints can be explored in a strictly data-driven and hypothesis-free manner to address the question posed above. Results gained from this research could be of particular interest for future studies in this field and also help to navigate through already existing inconsistencies in the literature.

## 2 Methods

### 2.1 Study overview and procedure

As part of a larger longitudinal study on app assisted learning, two age groups were recruited to participate in consecutive experimental sessions involving multiple data collection timepoints. These two studies included a group of children in one study and a group of adults in the other study, with age as a between-group variable. Both groups underwent various cognitive and neurophysiological tests over time and were required to use an app to learn new vocabulary in a second-language (L2) during the intervals between testing sessions, forming an overall mixed design.

Participants who fully completed two vocabulary tests, two resting state EEG recordings, and used the app between these sessions were included in the test-retest reliability analysis for the present two studies. Between the two timepoints (T1 and T2) for vocabulary tests and resting-state EEG sessions, participants used the app to study a second language. Consequently, both vocabulary tests and resting-state EEG sessions were scheduled to measure behavioral performance and neurophysiological changes before (T1) and after (T2) the language training with the app. A progress in language training and induced changes in LTM performance were anticipated during this time interval. The children group and the adult group were treated as separate studies. EEG power values and microstate measures were the dependent variables.

### 2.2 Study 1

#### 2.2.1 Participants

All child participants were recruited in the German and French-speaking regions of Switzerland, with native languages being either French or German. To ensure the validity of the results, only healthy participants, free from any chronic medical conditions, psychological disorders, language or memory deficits, were included in the study. A total of 44 children took part in the study. Of these, 36 participants (18 females, 23 righthanded, age M = 11.22 years, SD = 1.16, range = 9-15) fully completed the two testing sessions without technical problems and with the adequate data quality required for the present reliability analysis. All participants’ parents signed informed consent (study approved by UniDistance Suisse ethics committee; approval nr.: 2019-12-00002).

#### 2.2.2 Materials & Procedure

We employed a one-factorial within-subjects design with time (pre [T1]/post [T2] intervention) as the single factor. The dependent variables included performance from the vocabulary test, EEG power within the standard frequency bands, as well as variables derived from microstate analysis. This design allowed us to examine changes in both behavior and neurophysiological activity across the two time points, providing insights into the effects of the intervention (app). Each child initially completed a vocabulary test (T1), typically administered in a paper-and-pencil format within a classroom setting. This was followed by an EEG recording session (T1), conducted either on the same day or the following day. After completing T1, children used a language-learning app to study their second language. Participants were either native German or French speakers (L1) and used the app to learn vocabulary in their respective second language (L2), either French or German. Given that L2 is an official language and part of the standard school curriculum, children were exposed to L2 not only through the app but also through regular classroom instruction and additional resources such as textbooks during the study period. The vocabulary tests were tailored accordingly, assessing French or German vocabulary depending on each child’s L1. The frequency and intensity of app usage were left to the discretion of the individual participants. Following the app intervention, the children completed a second vocabulary test and EEG session (T2).

The vocabulary test was a cued-recall translation task comprising 20 items. Half of the items required translation from L1 to L2, while the other half required translation from L2 to L1. Participants provided their responses in written form. Responses on the translation items within a vocabulary test were marked as either correct or incorrect and the result was transformed into a percentage score representing behavioral performance. This design allows for the calculation of difference scores (T2 minus T1), representing an approximation to changes in LTM performance. Regardless of the individual app usage and language aptitude, each participant’s vocabulary test difference score (T2 minus T1) can be related to neurophysiological parameters, that show high reliability in our analysis. The percentage scores representing improvement in L2 were used as a proxy for LTM performance. These scores were correlated with EEG resting state measures at T1. This analysis aimed to estimate the predictive value of the stable neurophysiological markers at T1 for changes in LTM performance over time, as evidenced by the improvement in behavioral performance in L2 within the vocabulary test.

Participants sat in a chair with their eyes open and were told to fixate on a cross on a screen while the resting state EEG was recorded for 180 seconds. The 180-second-long eyes-open segments were pre-processed with the EEGLAB toolbox (v2023.0) for MATLAB (R2020b). The data was re-referenced to the common average and was filtered with a high-pass (0.5 Hz) and a low-pass (80 Hz) filter. Line-noise was removed at 50 Hz and its harmonics up to 150 Hz using the function ‘pop_cleanline()’ with default settings. Artifact rejection and removal of bad channels was performed using the function ‘pop_clean_rawdata()’ with standard parameters. Eye-movement artifacts and muscle artifacts were removed using Independent Component Analysis (ICA). ICA was performed on a copy of the data that was bandpass-filtered between 1-30 Hz and was then down-sampled to 100 Hz. Artifact-laden components were identified using ‘pop_iclabel()’ (Pion-Tonachini et al., 2019). All components related to eye-movements and muscle artifacts were removed from the original data. Finally, previously rejected channels were recovered using spherical interpolation, and all channels were again re-referenced to the common average.

#### 2.2.3 Data Analysis

The data were analysed in a data-driven, hypothesis-free manner. This approach allows for an unbiased examination of the stability of the different measures. In general data points outside 2 SD of the mean were removed before performing statistical tests. In all correlational analyses throughout the present studies, Bonferroni correction was applied to the p-values for all the correlations conducted to account for multiple (21) comparisons. All statistical analyses were conducted using R version 4.3.1 (R Core Team, 2023).

##### 2.2.3.1 EEG measures

###### 2.2.3.1.1 EEG Power

The EEGLAB function ‘spectopo()’ was used to obtain the average power spectral density (PSD) for 2-second epochs with an 80% overlap over the entire length of resting-state recording (180 seconds), ensuring a detailed frequency resolution. This analysis provided a frequency resolution of 0.5 Hz, covering a frequency range of 0.5 Hz to 80 Hz. The output from ‘spectopo()’ was converted from decibels (dB) to a linear scale (µV²/Hz) to facilitate further analysis. For each electrode, the absolute power within five standard frequency bands (delta: 1-4 Hz, theta: 4-8 Hz, alpha: 8-14 Hz, beta: 14-30 Hz, gamma: 30-75 Hz) was calculated over the specified frequency range. For each frequency band, the corresponding indices in the frequency vector were identified. The power within the band was then calculated by taking the integral of the PSD over the specified frequency range using the trapezoidal rule, which provides an estimate of the area under the curve of the power spectrum within that band. This process effectively quantifies the total power contained in each frequency band of the EEG signal.

Additionally, relative power for each frequency band was determined by normalizing the absolute power in each band to the total power across all frequencies resulting in a percentage value. Finally, to obtain a comprehensive measure of brain activity, the absolute and relative powers were averaged across all electrodes for each frequency band. This approach provided an overall representation of the brain’s spectral power distribution. For subsequent statistical analysis, the absolute power values for each frequency band were log-transformed using the base-10 logarithm. This transformation was performed to normalize the data distribution and stabilize variance, which are prerequisites for parametric statistical tests.

###### 2.2.3.1.2 Microstates

To calculate microstates, time points of interest for the analysis are typically identified using peaks in global field power (GFP) which represent pronounced EEG scalp topography. Such quasi-stable states of the electric field are assumed to last for short moments (60-120 ms) and presumably relate to differentiable functional states of the brain (Wackermann et al., 1993). Microstate analyses were done using the EEGLAB plugin MICROSTATELAB by Koenig (Nagabhushan Kalburgi et al., 2023). The ‘pop_FindMSMaps()’ function was used on the pre-processed EEG files to identify individual template maps (4-7 class number solutions in the k-means algorithm). Polarity was ignored, 20 restarts were used per dataset, and the clustering was completed on all GFP peaks. Four mean microstate maps across a single group within a time point (e.g. children T1) were identified using the ‘pop_CombMSMaps()’ function and four grand mean maps were identified across all means while ignoring polarity (Koenig et al., 2002). The solution of four classes of grand mean microstate maps was used to backfit the to the individual EEG data sets with the function ‘pop_FitMSMaps()’ as four maps are considered an optimal number for such analyses (Michel & Koenig, 2018). The mean duration, occurrence, coverage (fraction of total recording time that the microstate is dominant) and mean GFP of the four classes were calculated at the subject level and used for further statistical analysis.

###### 2.2.3.1.3 Reliability analysis

To measure the reliability of the EEG resting state measures between sessions, test-retest reliability was estimated using T1 and T2 measures. Interclass Correlation Coefficients (ICC) (Fisher, 1970) were calculated to quantify reliability, utilizing the R-package ‘psych’ (Revelle, 2023). ICC values range from 0 to 1, with higher values indicating greater reliability. Specifically, values from 0.5-0.75 indicate moderate reliability, values from 0.76-0.9 indicate good reliability, and values above 0.9 represent excellent reliability (Koo & Li, 2016). Significance tests further determine if the ICC is significantly different from 0 (alternative hypothesis).

To estimate the within-session stability of the relevant EEG resting state measures, split-half reliability measures were calculated. The 180-second-long raw segments from both T1 and T2 were divided into two 90 second segments each. Test-retest reliability was assessed using ICC values derived from two 180-second segments, while split-half reliability was calculated using the 90-second segments. The measures from the two 90-second segments within each timepoint (T1 and T2) were correlated using Spearman-Brown-Correction (Lopez et al., 2023). This method provides an additional estimate of split-half reliability for both T1 and T2, complementing the test-retest reliability of all relevant resting state measures. A measure was considered sufficiently reliable if both the split-half and test-retest reliability were equal to or exceeded .75.

### 2.3 Study 2

Unless otherwise specified, study 2 was conducted in the same manner as study 1.

#### 2.3.1 Participants

Out of 96 recruited participants, a total of 90 adults (58 females, 84 righthanded, age M = 28.96, SD = 6.85, range = 18-40) completed all required testing sessions without issues. All participants gave informed consent (study approved by UniDistance Suisse ethics committee; approval nr.: 2019-12-00002). Due to organizational and logistic reasons, vocabulary tests and resting-state EEG sessions could not always be scheduled on the same day for the adults. Some participants therefore completed T1 vocabulary and EEG recording in more than one session before starting the app usage.

#### 2.3.2 Material & Procedure

Adults used the app to practice Finish as a second language. The vocabulary tests were designed accordingly, with adults being tested on Finish material, with more items (48) and with computerized versions of the vocabulary test. Adults used the app following a more structured and consistent schedule, either for 1 hour per day over two weeks or approximately 3 hours every third day over three weeks.

## 3 Results

### 3.1 Study 1

Children took the vocabulary tests on average 71.33 days (SD = 26.12) apart and completed the EEG resting state recordings 59.69 days (SD = 22.08) apart.

#### 3.1.1 Behavioral measures

Figure 1 shows the results from the vocabulary test. At T1, children performed with an average score of 14.51% (SD = 11.04%) and improved to 42.16% (SD = 16.90%) at T2 after training with the app. The children’s test scores were significantly above zero at T1, as they already had some prior familiarity with the language. The difference score (T2 minus T1) of 27.65% (SD = 17.49%) represents the changes in LTM performance based on improvement in L2 within the children sample. A paired-samples t-test showed a significant difference between T1 and T2 *t*(33) = −9.98, *p* < .001, *d* = 2.07.

**Figure 1.**
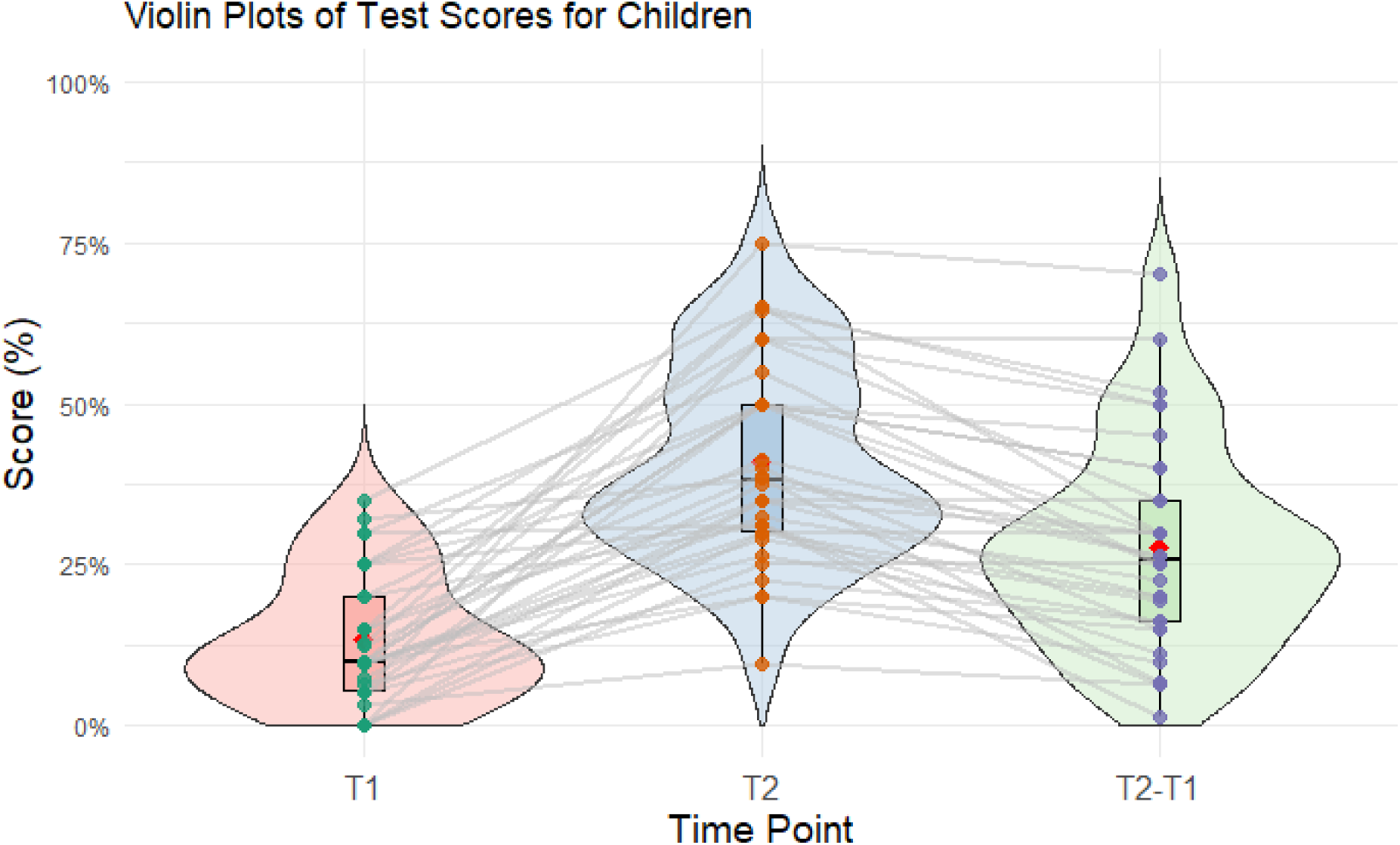
Violin plots for the performance on the vocabulary test in the children sample of study 1. The plots show the mean (red), the median (horizontal line), the boxplot, the interquartile range (vertical line) and the density distribution of the data at each timepoint (T1, T2 and T2 minus T1).

#### 3.1.2 EEG measures

##### 3.1.2.1 EEG Power

Table 1 shows the ICC results for the children sample in our study 1 for both absolute and relative power calculations. The ICCs from our study are listed next to ICCs reported in the literature (Lopez et al., 2023) based on similar calculations of absolute (Webb et al., 2023, but see Metzen et al., 2022 with higher ICC of 0.75 for absolute alpha power) and relative EEG power (Suárez-Revelo et al., 2015) on the whole brain during resting state recordings with healthy subjects. While all ICC values in the present study were significantly greater than 0 at the 5% alpha level only the bold numbers represent good reliability according to guidelines (ICC >= 0.75). The overall high split-half reliability calculations are also listed separately for T1 and T2.

**Table 1.** EEG Power Reliabilities.

| Power Measure | Test-Retest Reliability |  | Split-Half Reliability |  |
| --- | --- | --- | --- | --- |
|  | ICC present study | ICC | T1 | T2 |
|  | Children Study 1 | Literature | Children Study 1 |  |
| Absolute |  |  |  |  |
| Delta | 0.77 | 0.39 | 0.96 | 0.97 |
| Theta | 0.75 | 0.51 | 0.97 | 0.94 |
| Alpha | 0.81 | 0.67 | 0.94 | 0.94 |
| Beta | 0.76 | 0.68 | 0.98 | 0.94 |
| Gamma | 0.54 | 0.45 | 0.87 | 0.88 |
| Relative |  |  |  |  |
| Delta | 0.81 | 0.46 | 0.89 | 0.91 |
| Theta | 0.61 | 0.76 | 0.89 | 0.94 |
| Alpha | 0.84 | 0.75 | 0.92 | 0.96 |
| Beta | 0.80 | 0.63 | 0.97 | 0.98 |
| Gamma | 0.59 | 0.57 | 0.88 | 0.93 |

*Note.* ICCs from the literature are based on Lopez et al. (2023), bold ICC values represent good test-retest reliability (ICC $\geq 0.75$ ) in the present study. Split-Half Reliability was calculated with the Spearman-Brown-Correction (between the two halves of 90 s segments). n. sig. $p > .05$

##### 3.1.2.2 Microstates

Figure 2 shows the four grand mean microstate maps when data from both studies are combined. Table 2 displays the reliability estimates for the microstate measurements (mean GFP, duration, occurrence, duration and coverage of the four classes) in the children sample of study 1. Split-half reliability and ICC values are overall smaller for microstate measures than for EEG power. Except for one ICC value, all microstate measures in the present study showed an ICC significantly greater than 0 at the 5% alpha level, similar to other studies (Popov et al., 2023).

**Figure 2.**
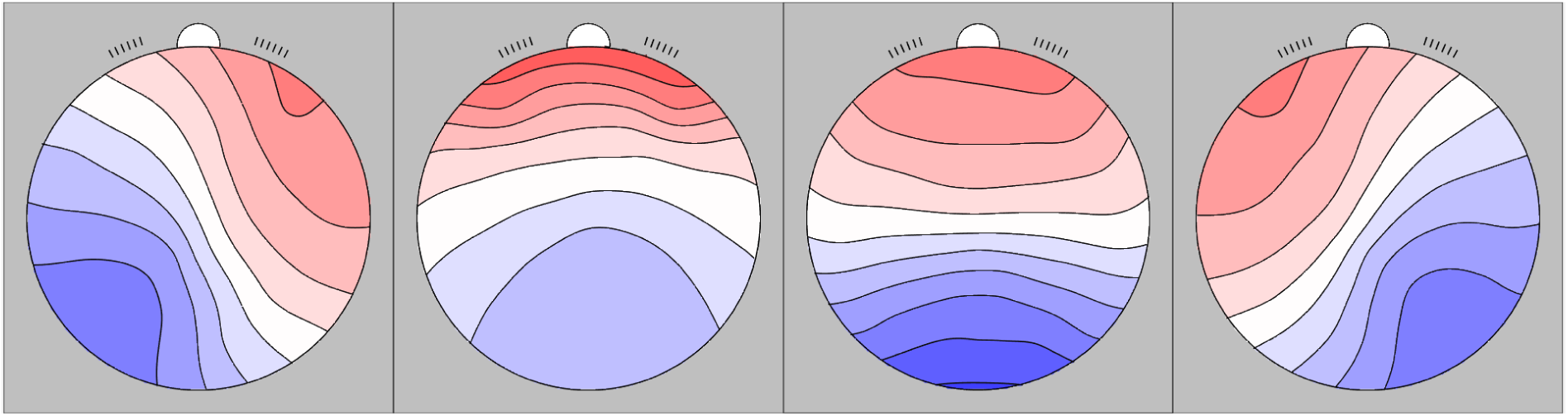
Grand mean microstate maps (A, B, C, D) identified in the present study.

**Table 2.**
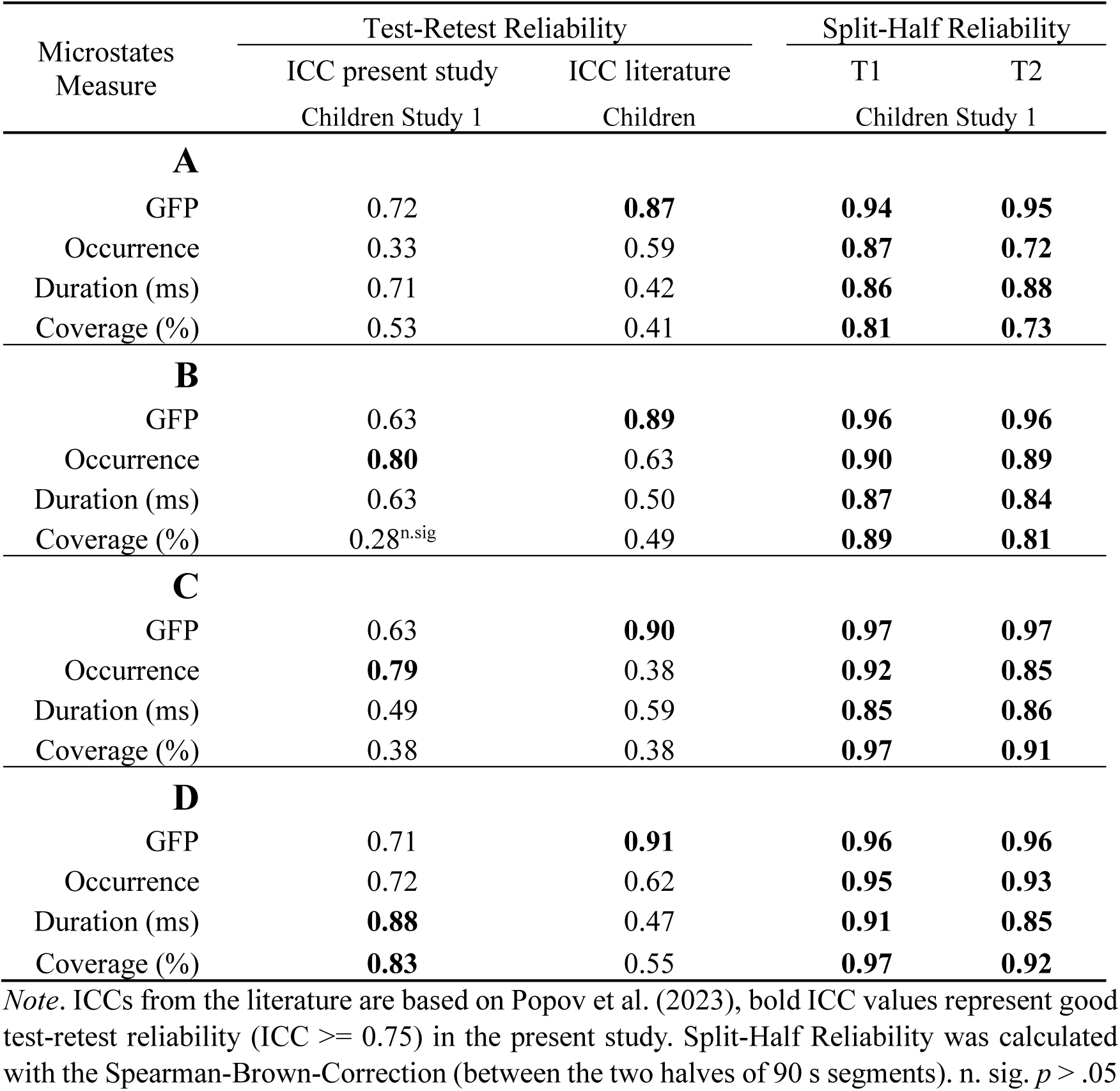
EEG Microstates Reliabilities.

#### 3.1.3 Correlational analyses

Focusing only on sufficiently reliable power measures (both test-retest and split-half reliability >= 0.75) all correlations (Pearson) were calculated between the reliable power measures at T1 (7 measures of EEG power) and the change in LTM performance between T1 and T2 in the children sample of study 1. The results (Bonferroni corrected) are shown in table 3. There was no significant correlation in this analysis.

**Table 3.** Correlations between reliable EEG power measures and LTM.

|  |  |  |  |  |
| --- | --- | --- | --- | --- |
| **Absolute** | Delta | Theta | Alpha | Beta |
| LTM | 0.19 | 0.16 | -0.15 | 0.08 |
| **Relative** | Delta | - | Alpha | Beta |
| LTM | 0.25 | - | -0.36 | -0.05 |

Focusing only on sufficiently reliable microstate measures (both test-retest and split-half reliability >= 0.75) all correlations (Pearson) were calculated between the reliable power measures at T1 (4 measures of EEG power) and the change in LTM performance between T1 and T2 in the children sample of study 1. The results (Bonferroni corrected) are shown in table 4. There was no significant correlation in this analysis.

**Table 4.**
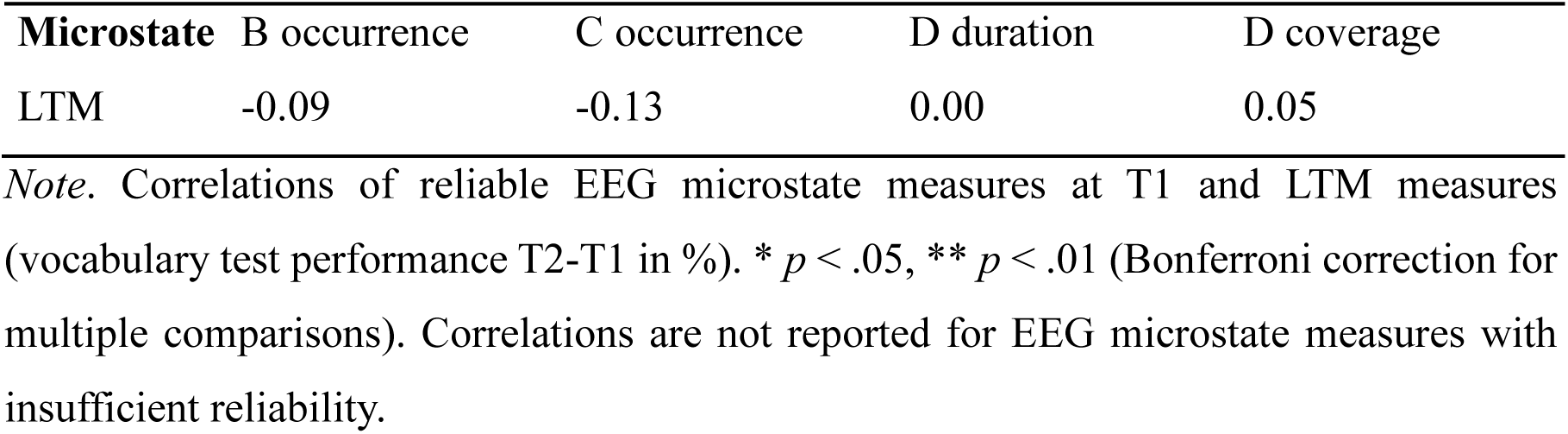
Correlations between reliable EEG microstates measures and LTM.

| **Microstate** | **B occurrence** | **C occurrence** | **D duration** | **D coverage** |
| --- | --- | --- | --- | --- |
| LTM | -0.09 | -0.13 | 0.00 | 0.05 |

*Note.* Correlations of reliable EEG microstate measures at T1 and LTM measures (vocabulary test performance T2-T1 in %). \* $p < .05$ , \*\* $p < .01$ (Bonferroni correction for multiple comparisons). Correlations are not reported for EEG microstate measures with insufficient reliability.

### 3.2 Study 2

In the adult sample, the two sessions with the vocabulary test were on average 29.86 days (SD = 28.09) apart and 14.13 days (SD = 0.77) elapsed between the first and the second EEG resting state recordings.

#### 3.2.1 Behavioral measures

Adults showed a significant improvement in LTM performance from T1 to T2. This was indicated by a change in retrieval performance of 65.13% (SD = 21.75%) with an average of 0.00% (SD = 0.50%) at T1 and an average of 65.13% (SD = 21.76%) at T2. A paired-samples t-test showed a significant difference between T1 and T2 *t*(82) = −32.66, *p* < .001, *d* = 5.06.

#### 3.2.2 EEG measures

##### 3.2.2.1 EEG Power

Table 5 shows the ICC results for the adult sample in our study 2 both for absolute and relative power calculations. Overall, the split-half reliabilities in this sample were very high and the highest ICC values were observed in the alpha band.

**Figure 3.**
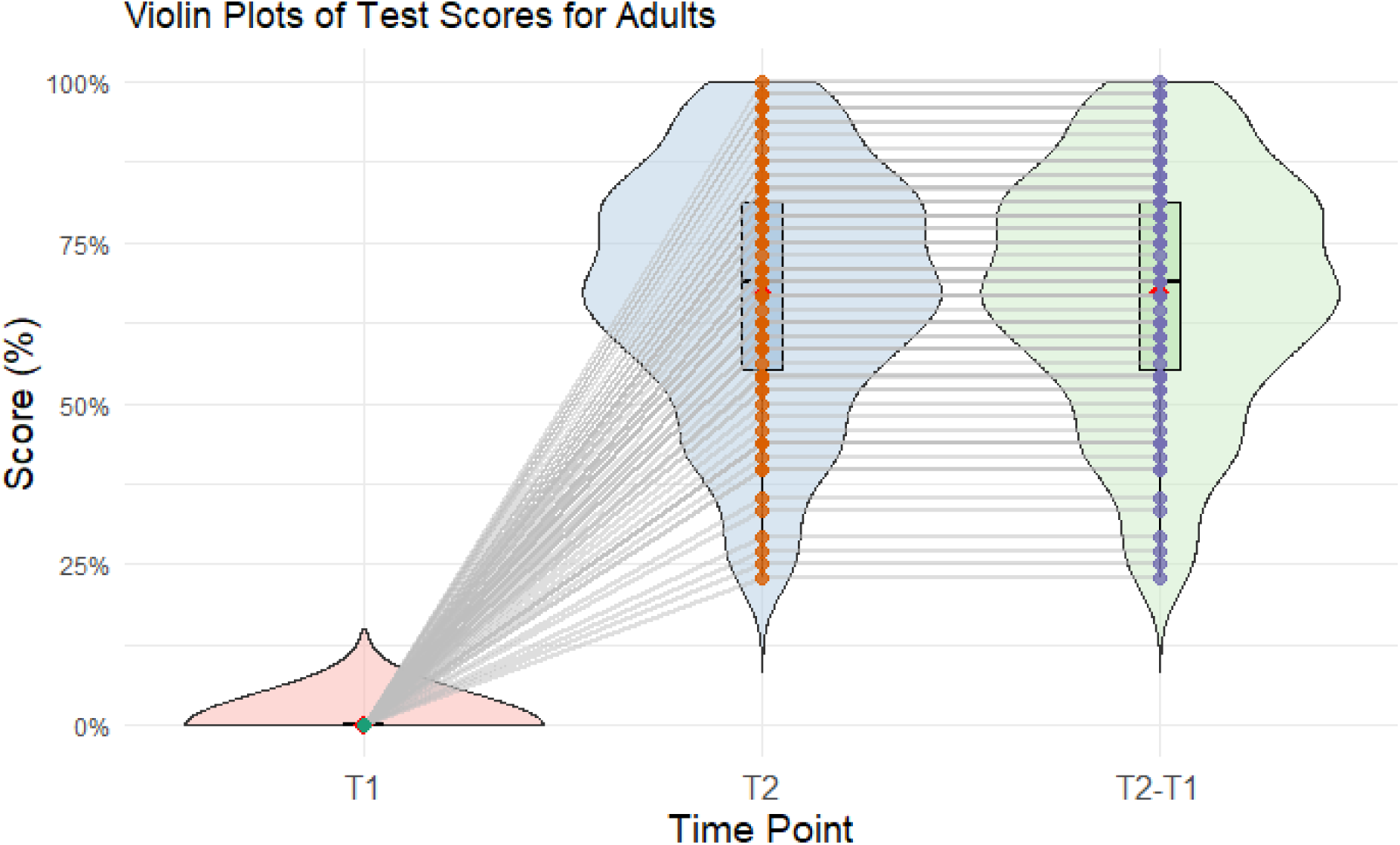
Violin plots for the performance on the vocabulary test in the adult sample of study 2. The plots show the mean (red), the median (horizontal line), the boxplot, the interquartile range (vertical line) and the density distribution of the data at each timepoint (T1, T2 and T2 minus T1).

**Table 5.**
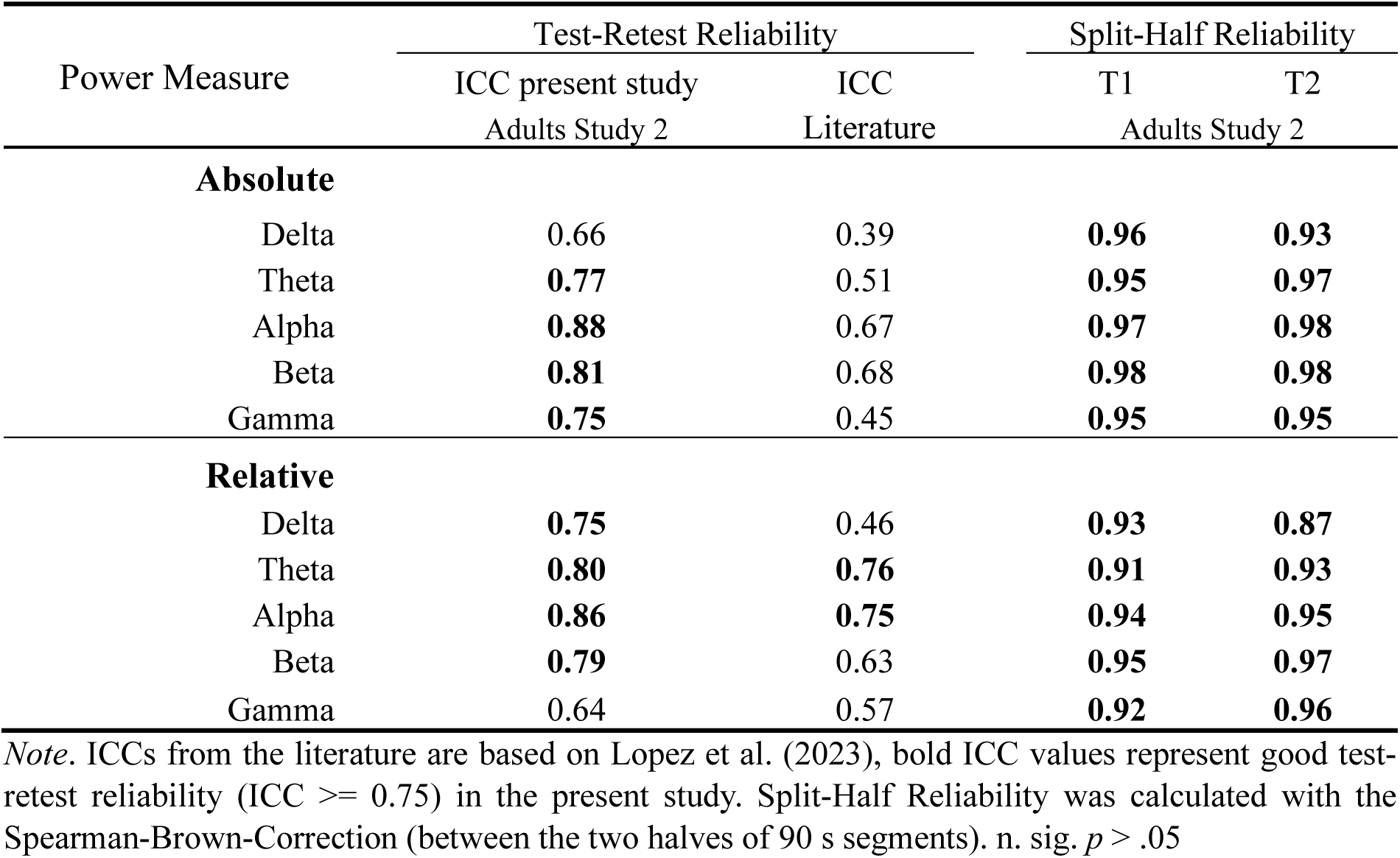
EEG Power Reliabilities.

| Power Measure | Test-Retest Reliability |  | Split-Half Reliability |  |
| --- | --- | --- | --- | --- |
|  | ICC present study | ICC | T1 | T2 |
|  | Adults Study 2 | Literature | Adults Study 2 |  |
| Absolute |  |  |  |  |
| Delta | 0.66 | 0.39 | 0.96 | 0.93 |
| Theta | 0.77 | 0.51 | 0.95 | 0.97 |
| Alpha | 0.88 | 0.67 | 0.97 | 0.98 |
| Beta | 0.81 | 0.68 | 0.98 | 0.98 |
| Gamma | 0.75 | 0.45 | 0.95 | 0.95 |
| Relative |  |  |  |  |
| Delta | 0.75 | 0.46 | 0.93 | 0.87 |
| Theta | 0.80 | 0.76 | 0.91 | 0.93 |
| Alpha | 0.86 | 0.75 | 0.94 | 0.95 |
| Beta | 0.79 | 0.63 | 0.95 | 0.97 |
| Gamma | 0.64 | 0.57 | 0.92 | 0.96 |

Note. ICCs from the literature are based on Lopez et al. (2023), bold ICC values represent good test-retest reliability (ICC >= 0.75) in the present study. Split-Half Reliability was calculated with the Spearman-Brown-Correction (between the two halves of 90 s segments). n. sig. $p > .05$

##### 3.2.2.2 Microstates

Table 6 displays the reliability estimates for the microstate measurements (mean GFP, duration, occurrence, duration and coverage of the four classes) in the adult sample of study 2. Split-half reliabilities and ICC values are overall smaller for microstate measures than for EEG power. All microstate measures in the present study showed an ICC significantly greater than 0 the 5% alpha level, similar to other studies (Popov et al., 2023). However, only two ICC values (bold) represent good reliability according to guidelines.

**Table 6.** EEG Microstates Reliabilities.

| Microstates Measure | Test-Retest Reliability |  | Split-Half Reliability |  |
| --- | --- | --- | --- | --- |
|  | ICC present study | ICC literature | T1 | T2 |
|  | Adults Study 2 | Adults | Adults Study 2 |  |
| **A** |  |  |  |  |
| GFP | 0.71 | **0.78** | **0.95** | **0.96** |
| Occurrence | 0.66 | 0.06 | **0.85** | **0.89** |
| Duration (ms) | 0.68 | 0.70 | **0.86** | **0.92** |
| Coverage (%) | 0.55 | 0.22 | **0.81** | **0.90** |
| **B** |  |  |  |  |
| GFP | 0.71 | 0.69 | **0.97** | **0.97** |
| Occurrence | 0.57 | 0.39 | **0.87** | **0.95** |
| Duration (ms) | 0.63 | 0.09 | **0.86** | **0.93** |
| Coverage (%) | 0.38 | 0.28 | **0.85** | **0.92** |
| **C** |  |  |  |  |
| GFP | 0.66 | 0.70 | **0.94** | **0.96** |
| Occurrence | 0.68 | 0.51 | **0.92** | **0.95** |
| Duration (ms) | 0.59 | 0.17 | **0.79** | **0.93** |
| Coverage (%) | 0.59 | 0.33 | **0.84** | **0.91** |
| **D** |  |  |  |  |
| GFP | 0.71 | **0.75** | **0.97** | **0.97** |
| Occurrence | 0.74 | 0.35 | **0.92** | **0.95** |
| Duration (ms) | **0.80** | 0.39 | **0.93** | **0.94** |
| Coverage (%) | **0.79** | 0.21 | **0.96** | **0.95** |

Note. ICCs from the literature are based on Popov et al. (2023), bold ICC values represent good test-retest reliability (ICC $\geq 0.75$ ) in the present study. Split-Half Reliability was calculated with the Spearman-Brown-Correction (between the two halves of 90 s segments). n. sig. $p > .05$

#### 3.2.3 Correlational analyses

Focusing only on sufficiently reliable power measures (>= 0.75) all correlations (Pearson) were calculated between the reliable power measures at T1 (8 measures of EEG power) and the LTM in the adult sample of study 2. The results are shown in table 7. Alpha power values were significantly positively correlated with LTM performance in the adult sample. A scatterplot for the significant correlations are depicted in figure 4 and 5.

**Figure 4.**
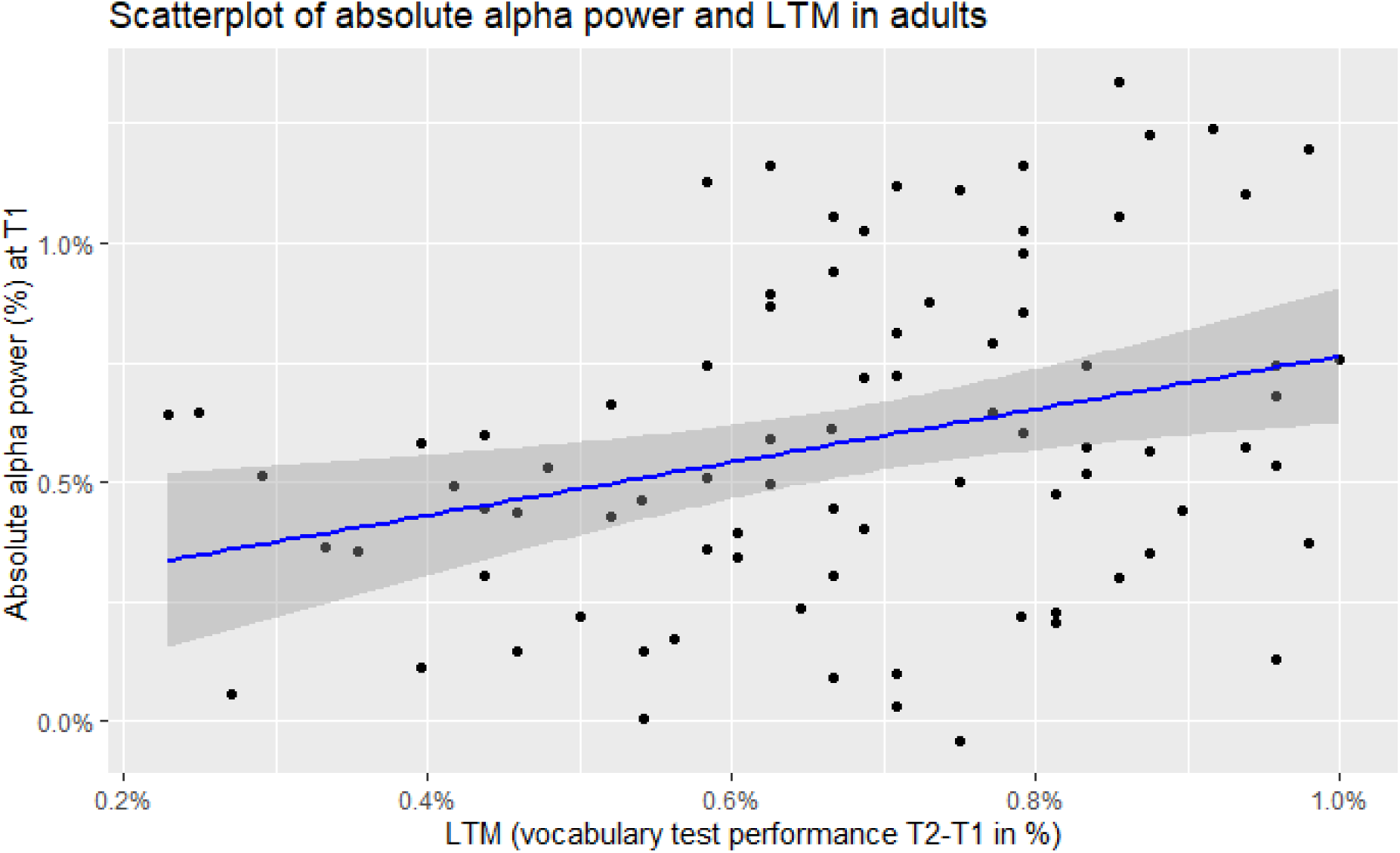
Scatterplot of absolute alpha power (%) and LTM (vocabulary test performance T2-T1 in %) for the adult sample of study 2.

**Figure 5.**
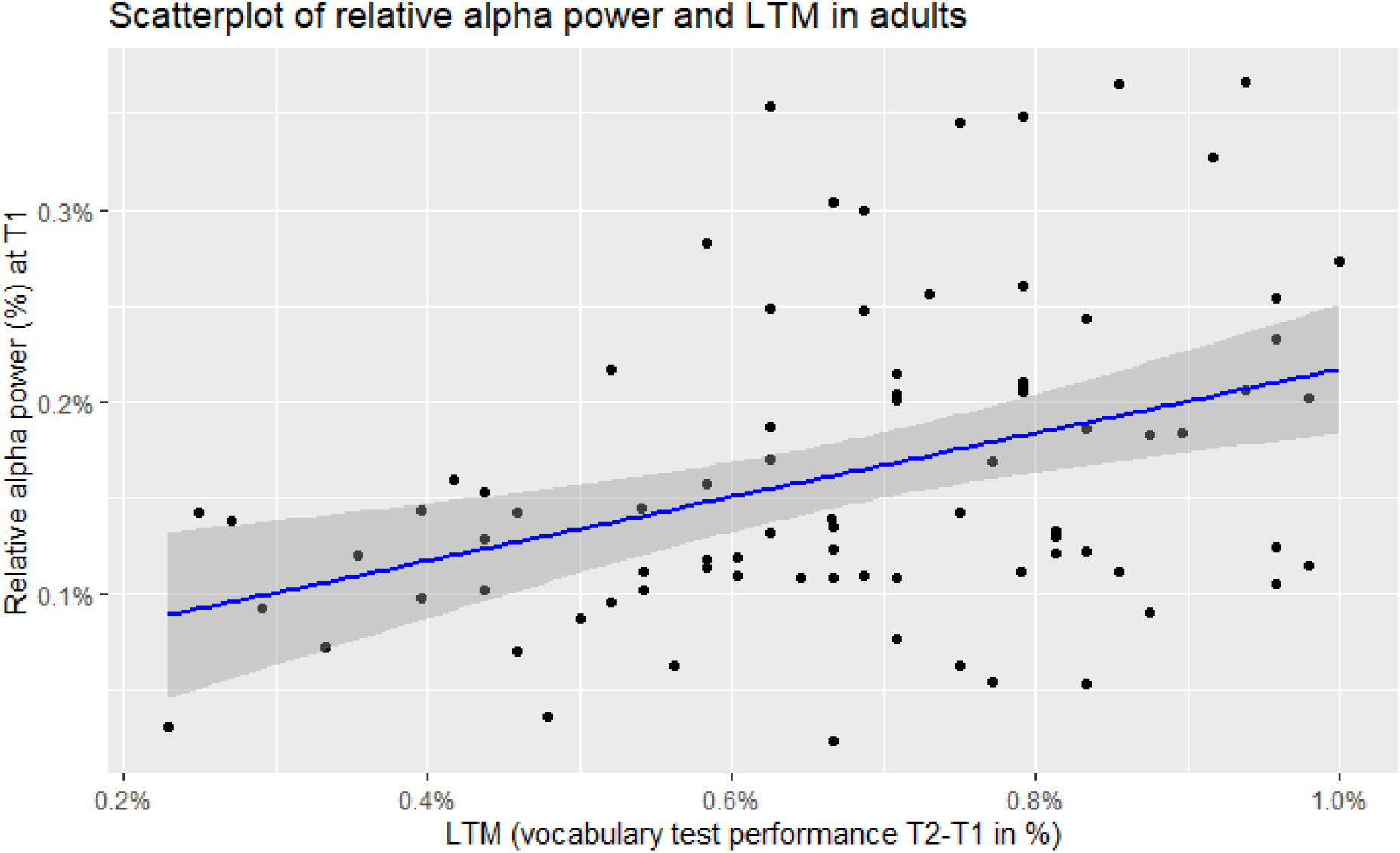
Scatterplot of relative alpha power (%) and LTM (vocabulary test performance T2-T1 in %) for the adult sample of study 2.

**Table 7.** Correlations between reliable EEG power measures and LTM.

| **Absolute** | - | Theta | Alpha | Beta | Gamma |
| --- | --- | --- | --- | --- | --- |
| LTM | - | 0.07 | 0.31* | -0.09 | -0.24 |
| **Relative** | Delta | Theta | Alpha | Beta | - |
| LTM | 0.02 | 0.12 | 0.38** | 0.05 | - |

*Note.* Correlations of reliable EEG power measures at T1 and LTM measures (vocabulary test performance T2- T1 in %). \* $p < .05$ , \*\* $p < .01$ (Bonferroni correction for multiple comparisons). Correlations are not reported for EEG power measures with insufficient reliability.

Again, focusing only on sufficiently reliable microstate measures (>= 0.75) all correlations (Pearson) were calculated between the reliable microstate measures at T1 (2 microstate measures) and the LTM performance in the adult sample of study 2. The results are shown in table 8. No correlations were significant.

**Table 8.** Correlations between reliable EEG microstates measures and LTM.

| **Microstate** | D duration | D coverage |
| --- | --- | --- |
| LTM | 0.18 | 0.07 |

*Note.* Correlations of reliable EEG microstate measures at T1 and LTM measures (vocabulary test performance T2-T1 in %). \* $p < .05$ , \*\* $p < .01$ (Bonferroni correction for multiple comparisons). Correlations are not reported for EEG microstate measures with insufficient reliability.

## 4 Discussion

Both children and adults experienced a change in LTM performance over time as measured in successful L2 learning. An overall high split-half reliability was observed for almost all measures derived from resting-state EEG. More measures of EEG power resulted in high test-retest reliability as compared to measures of microstates. Furthermore, only a few of these measures were associated with LTM performance after correcting for multiple comparisons. Higher absolute and relative alpha power at T1 was the only EEG signal positively related to a change in LTM performance from T1 to T2 in our sample and this was only observed for the adult participants. Increased alpha power at the scalp could be a reliable neurophysiological marker in the adult brain for LTM formation, resulting in observable behavioral outcomes. Increased alpha power is assumed to facilitate memory processes by suppressing unwanted or irrelevant memory traces (Hanslmayr et al., 2012) and surface EEG could be a reliable tool for capturing this specific neural mechanism.

The high stability of the alpha band compared to other frequencies has also been noted in other studies (Lopez et al., 2023; Popov et al., 2023). Our results further suggest that absolute alpha power is more reliably related to changes in LTM performance than its proportion relative to the total power across all frequency bands.

Alpha power at the scalp in general can be interpreted as a modulation by thalamo-cortical networks (Bazanova & Vernon, 2014). Alpha power has a high heritability (Smit et al., 2006), is related to higher memory performance (Klimesch, 1999) and shows predictive power for language processing at the subject level (Bornkessel-Schlesewsky et al., 2022). Additionally, event-related alpha power has also been linked to lexical retrieval (Röhm et al., 2001; for a review see Bastiaansen & Hagoort, 2006; Rossi et al., 2023) and progress in L2 learning (Allal-sumoto & Duygu, 2024). While alpha power at the scalp appears to be related to LTM, intracranial recordings with relation to LTM consolidation have reported findings in the gamma range (Axmacher et al., 2008). Other studies related intracranial theta to subsequent memory (for a review see Johnson et al., 2015). LTM relevant brain mechanisms could manifest themselves in different frequencies at the scalp compared to intracranial level. This difference could be due to the intracranial recordings being conducted in the Hippocampus. Such oscillations are less likely to be picked up by electrodes on the scalp. One study demonstrated a correspondence between intracranial and scalp EEG during successful memory encoding (Long et al., 2014). Specifically, decreases in lower frequencies (theta: 3–8 Hz) and increases in higher frequencies (gamma: 44–100 Hz) were observed to accompany recalled items in time-frequency spectrograms. However, no significant correspondence between intracranial and scalp EEG was reported for the alpha range (10–14 Hz).

There is a difference between the children and adult sample in the present study. Different neuronal mechanisms could guide changes in LTM performance through L2 learning in children and adults. While lower occipital alpha power (7-10 Hz) could be related to enhanced sensory processes through inhibited neural circuits in children (Kwok et al., 2019), higher left and frontal alpha power (10-13 Hz) could be related to enhanced retrieval of semantic information from LTM in adults (Grabner et al., 2007). Such differences between the age-groups could explain the lack of significant correlation in the children sample.

The results in the present two studies could explain some inconsistent findings in the literature when comparing power measures and LTM based on L2 acquisition. As shown here, future studies could benefit from focusing on stable power measures e.g. alpha power and generally relying on relative power measures. While the results in alpha power seem promising and alpha power measures are considered highly important in understanding human cognition, it is crucial to note, that this power measure is not at all specific for LTM and has been associated with a wide variety of cognitive functions and impairments (Başar & Güntekin, 2012). LTM was the focus of the present study but any underlying cognitive process resulting in proficiency at second language acquisition could also be reflected in pronounced alpha power. While the reliability and stability of this measure seem robust, future studies should try to exclude alternative cognitive processes distinguishable from LTM as explanations for the effect in alpha power.

While the present results showed moderate to good reliability for some microstate measures, their relation to LTM performance appears to be nonsignificant in our samples. Microstates have sometimes been used to investigate language processes (Koenig & Lehmann, 1996) but this was mostly done with experimental tasks focusing on event-related brain potentials (Brandeis et al., 1995; Koenig et al., 1998) and less in resting-state recordings. When using lexical tasks and event-related microstates, high stability and reliability have also been reported (Laganaro, 2017), and microstates have been linked to language capabilities (Jouen et al., 2021; Tanaka et al., 2002). Such studies, however, focused less on memory and more on short-term language-specific processes e.g. speech production. Only on rare occasions were event-related microstates associated with language related stimuli (e.g. syllables) with implications on long-term cognitive processes (e.g. Giroud et al., 2017). In one such study second language acquisition over a period of 5 months was related to a shortened microstate presumably representing an improved access to the semantic network and faster as well as more automated processing of L2 words (Stein et al., 2006). Contrary to our study these microstates were calculated in an event-related lexical task. These results were related to shorter activation of the inferior frontal gyrus (left). Both studies relied on a multitude of instructions and tasks with detailed trial-sequences showing the pitfalls of ERP studies outlined above without being replicated but were able to directly relate LTM relevant stimulation to metrics from their microstate analyses.

In the present studies, microstates were calculated from resting-state recordings. As in a previous study (Antonova et al., 2022), microstate duration showed somewhat higher reliability than occurrence. However, the relation of resting-state microstates to LTM in the literature is less clear. A recent registered report with 140 healthy participants showed no significant correlation between any microstate measurements and a series of cognitive tests on executive function (e.g., letter memory task, dual N-back) (Chenot et al., 2024). Although LTM was not the focus here, executive functions rely on cognitive processes also relevant for LTM. This absence of any relation between microstate measures and cognitive performance in tasks of executive functions might also explain the absence of any such correlation in the present study. The relationship between microstates and behavioural indices of cognition could be more tangible in event-related tasks than in resting-state recordings. Another explanation could be that this relationship is more pronounced in clinical samples and not in healthy participants like in Chenot et al. (2024) or the present study. In a small-sampled study, Grieder et al. (2016) found a significant correlation between microstate measures and a language test in eight patients suffering from semantic dementia. Although when using the same language test, another study with 117 patients with Alzheimer’s disease and 117 patients with mild cognitive impairment, reported no significant correlation between the test scores and any microstate measure (Musaeus et al., 2020). In summary, microstates represent a very useful and easily employable analysis technique but their use in examining LTM in healthy subjects still needs further clarification. Recommendations on future directions have been pointed out (Khanna et al., 2015).

Some notable limitations in the present study must be emphasized. While the different intervals between T1 and T2 (vocabulary test and EEG) for the two age groups are less relevant, the different sample sizes are quite noticeable. With more than twice the sample size in the adult study, the generally more positive results for this age-group could be explained with a simple lack of statistical power in the children group. Another aspect is visualized in the behavioral results where LTM performance (T2 minus T1 in the vocabulary test) scores show a much wider distribution and high variability in children than in adults. Most adults seem to have similar changes in scores over time and a narrower distribution. This inconsistent performance in children could also be a reason why no EEG measure, despite good reliability, was significantly correlated with this behavioral measure in children. The non-perfect reliability of the LTM performance measures could have reduced our ability to find associations between EEG measures and LTM performance. Another reason for the different results in the two studies could be that children studied a somewhat familiar language while adults had to learn an entirely new language. In the children’s study, the second language tested (French/German) corresponds to one of the official national languages of Switzerland. Consequently, children likely had some prior exposure to and knowledge of this second language through their everyday environment or schooling. In contrast, in the adults’ study, the second language tested (Finnish) was entirely new to all participants, as Finnish is neither an official language of Switzerland nor a language to which participants had prior exposure. Even when resting-state recordings make it easier to study children with EEG than event-related tasks, the vocabulary testing sessions are still prone to commonly associated problems when doing research with children. While new methods for resting state EEG analysis are constantly being developed, sometimes specifically within language research (Roehm et al., 2004), there is also a need to identify sources for the heterogenous results in the existing literature and establish metrics of reliability within a field for it to move forward.

### 4.1 Conclusion

This study set out to test the stability of two very promising resting-state EEG methodologies by calculating both test-retest and split-half reliabilities. Based on the present results, power values produce mostly, and microstates produce some stable measures. Future inconsistent findings could be avoided by focusing more on the stable measures within these two EEG methodologies. Despite their stability, their exact relationship with LTM remains a challenging question. Out of all the identified stable measures only relative alpha power appears to be linked to LTM and only in adult participants. This finding, if replicated should be evaluated using different approximations to LTM than L2 vocabulary tests and a larger children sample.

## Data and Code Availability

All data and analysis scripts used in this study are publicly available at the Open Science Framework (OSF): https://osf.io/5znyk/, DOI: https://doi.org/10.17605/OSF.IO/5ZNYK.

## Author Contributions

Nicolas Rothen and Thomas P. Reber conceived and organized the study and secured funding. Anastasios Ziogas and Simon Ruch performed the data analyses and wrote the manuscript. Nicole H. Skieresz and Sandy C. Marca were responsible for participant recruitment and data collection. All authors reviewed and approved the final version of the manuscript.

